# Unveiling unique metabolomic and transcriptomic profiles in three Brassicaceae crops

**DOI:** 10.1101/2024.12.18.629212

**Authors:** Liyong Zhang, Isobel A. P. Parkin

## Abstract

*Brassica napus*, *Camelina sativa* and field pennycress (*Thlaspi arvense*), represent one highly economically valuable crop and two emerging oilseed crops of the Brassicaceae family, respectively. As sessile organisms, these crops are continuously exposed to various stresses when grown in the field. Interestingly, the responses of these three crops to different environmental stimuli vary to a great extent, but there is very limited knowledge about the molecular basis of these differential responses. In this study, we employed untargeted metabolomics to compare the metabolic profile of these crops, and examined the potentially related genes through further integration with transcriptomic analysis. Our data revealed distinctive overall metabolic profiles among these three crops, where in particular, a variety of phenylpropanoids showed differential accumulation and the corresponding putative genes’ expression varied significantly. The results provide a valuable resource for those studying Brassicaceae species and will provide insight into the understanding of metabolic variation among these three important oilseed crops, and provide potential targets for the future breeding of stress tolerant crops.

## Introduction

The Brassicaceae family contains many economically important species, which are widely used as sources of oil and food, as well as ornamental plants (Raza et al., 2020). For oilseed crops, the most productive cultivated species is *Brassica napus*, which is a hybrid species derived from an interspecific cross between *Brassica oleracea* and *Brassica rapa*, and is grown as both an annual and biennial crop, mainly for oil extraction, in many countries (Kirkegaard et al., 2021). More recently two lesser known oilseeds from the Brassicaceae family have been garnering interest. One is *Camelina sativa*, which has a versatile oil profile and a unique ability to adapt to many different environmental conditions, along with several other favourable agronomic traits, that has generated worldwide recognition of its potential as a crop (Berti et al., 2016). The second, field pennycress (*Thlaspi arvense*), has become known as an attractive non-food oilseed crop for biodiesel given its high content of monounsaturated fatty acids (Zanetti et al., 2019).

In their natural field environment, along with various abiotic stresses, these oilseed crops are exposed to the pathogens and insects common to crops in the Brassicaceae family. Interestingly, these three closely related oilseed species display tolerance variations towards different stresses. Such as, *B. napus* and pennycress are very susceptible to drought, while *C. sativa* shows a greater degree of tolerance (Gugel & Falk, 2006; Vollmann & Eynck, 2015). Flea beetles commonly feed on plants of the Brassicaceae family, causing serious damage to the young seedlings (Li et al., 2024), where *B. napus* is very susceptible to flea beetles, but *C. sativa* and pennycress show strong resistance (Soroka & Grenkow, 2013). With their sessile nature, plants have to adapt rapidly to unfavorable environmental conditions (i.e. abiotic and biotic stresses), and to successfully cope with the diverse environmental stimuli, plants have evolved a set of sophisticated strategies including complex physiological and metabolomic changes (Bartwal et al., 2013; Hanada et al., 2008; He et al., 2018). The plant metabolome consists of hundreds of thousands of organic compounds, which can be divided into two general groups: primary and secondary metabolites, whereat primary metabolites are essential for the normal plants’ growth and development, and secondary metabolites are important for plant survival by mediating plant-environment interactions under unfavorable conditions (Erb & Kliebenstein, 2020; Llanes et al., 2018; Salam et al., 2023).

One unique group of secondary metabolites that are found mainly in Brassicaceae plants are glucosinolates (GSLs), which are nitrogen and sulphur-containing compounds (Prieto et al., 2019). Based on the amino acid precursor, GSLs can be classified into three different types, i.e. aliphatic, aromatic and indole GSLs (Halkier & Gershenzon, 2006). GSLs have been documented to be involved in various responses to abiotic and/or biotic stresses within different members from the Brassicaceae family (Chhajed et al., 2020; Del Carmen Martinez-Ballesta et al., 2013; Variyar et al., 2014). Interestingly, *B. napus* and pennycress have been reported to contain high amounts of aliphatic GSLs in their leaves, on the contrary, *C. sativa* has almost no detectable GSLs present in the leaves (Czerniawski et al., 2021). Whether these differences in GSLs among these three oilseeds will affect their responses to various environmental perturbations are largely unknown. Beyond GSLs, little is known regarding the potential metabolites that differentiate these three species, with their varying responses to important abiotic and biotic stresses. An effective way to study the overall metabolic profile in plants is metabolomics, which consists of two general approaches: targeted and untargeted analyses (Martins et al., 2021). Untargeted metabolomics analysis has been successfully applied to large-scale metabolic profiling to identify discriminative metabolites between different plant species and/or in response to environmental stimuli (Allwood et al., 2021; Arbona et al., 2013; Castro-Moretti et al., 2020). More recently, the integration of metabolomics with transcriptomics provides a more comprehensive view of gene-metabolite pairs, which allow us to explore the correlation between the transcriptional and metabolic profiles (Arias et al., 2023; Wang et al., 2021; Y. Zhao et al., 2019).

In this study, we employed untargeted metabolomics and transcriptomics to analyze the metabolites of *B. napus*, *C. sativa*, and pennycress, our main objectives were to 1) provide a comprehensive overview of metabolites in the leaves and cotyledons of these three oilseed crops; 2) identify the discriminative compounds among these three species; 3) attempt to elucidate the potential links between metabolites and regulatory genes.

## Results

### 1, Summary of metabolomics data

To get a general idea of the metabolite profile of leaves and cotyledons from *B. napus*, *C. sativa* and *T. arvense* (field pennycress), we carried out untargeted metabolomics through LC-MS analysis on a Chemical Isotope Labeling (CIL) Metabolomics Platform (S. Zhao et al., 2019). In total, of the thousands of metabolites that were detected 718 could be classified with confidence. The metabolites were classified into the following categories: short and medium-chain fatty acids, polyamine, neurotransmitter, phytohormone, phenol and quinone, flavonoid, vitamins and derivatives, phenylpropanoid, polyphenol, lipid, amino acids and derivatives, dipeptides and tripeptides, alkaloid, terpene, and others (Table 1). The most abundant two categories are amino acids and derivatives, dipeptides and tripeptides, which contributed 131 (18.2%) and 122 (16.9%) compounds respectively.

**Table 1.** Classification of metabolites identified in untargeted metabolomics analysis.

| Categories | *C. sativa* | *B. napus*<br>(DH12075) | *B. napus*<br>(N99) | pennycress | Overall |
| --- | --- | --- | --- | --- | --- |
| Short & Medium-Chain Fatty Acids | 9 | 9 | 9 | 9 | 9 |
| Polyamine | 6 | 6 | 6 | 6 | 6 |
| Neurotransmitter | 5 | 5 | 7 | 5 | 7 |
| Phytohormone | 7 | 7 | 8 | 8 | 8 |
| Phenol and Quinone | 3 | 3 | 3 | 3 | 3 |
| Flavonoid | 14 | 15 | 14 | 13 | 15 |
| Vitamins & Derivatives | 6 | 6 | 6 | 6 | 6 |
| Phenylpropanoid | 39 | 39 | 40 | 37 | 40 |
| Polyphenol | 4 | 4 | 4 | 4 | 4 |
| Lipid | 7 | 7 | 7 | 7 | 7 |
| Amino Acids & Derivatives | 131 | 131 | 130 | 130 | 131 |
| Dipeptides & Tripeptides | 122 | 121 | 122 | 121 | 122 |
| Alkaloid | 31 | 32 | 32 | 32 | 32 |
| Terpene | 3 | 3 | 3 | 3 | 3 |
| Others | 316 | 322 | 323 | 323 | 325 |
| Total metabolites | 703 | 711 | 714 | 707 | 718 |

To have an overview of the metabolites’ distribution across all examined samples, we performed a hierarchical clustering analysis, the clustering results indicated that the 718 metabolites could be classified into 10 subclasses according to their relative abundance (Figure 1). The abundance level of metabolites in each subclass varied to a great extent across the experimental samples. Notably, subclasses 2, 3, and 7 exhibited distinctive patterns, which effectively distinguished *B. napus*, *C. sativa*, and pennycress from one another. Metabolites in subclass 2 displayed significantly higher levels of accumulation in both Camelina’s leaf and cotyledon tissues; metabolites in subclass 3 demonstrated higher levels in both *B. napus* lines compared to *C. sativa* and pennycress. In contrast, metabolites in subclass 7 showed a much greater accumulation in pennycress (Figure 2). Analyses of each of the three differential subclasses showed no significant enrichment for metabolites in any one biochemical pathway. The specific details regarding the 10 subclasses as well as their included metabolites can be found in Supplemental Data1.

**Fig 1.**
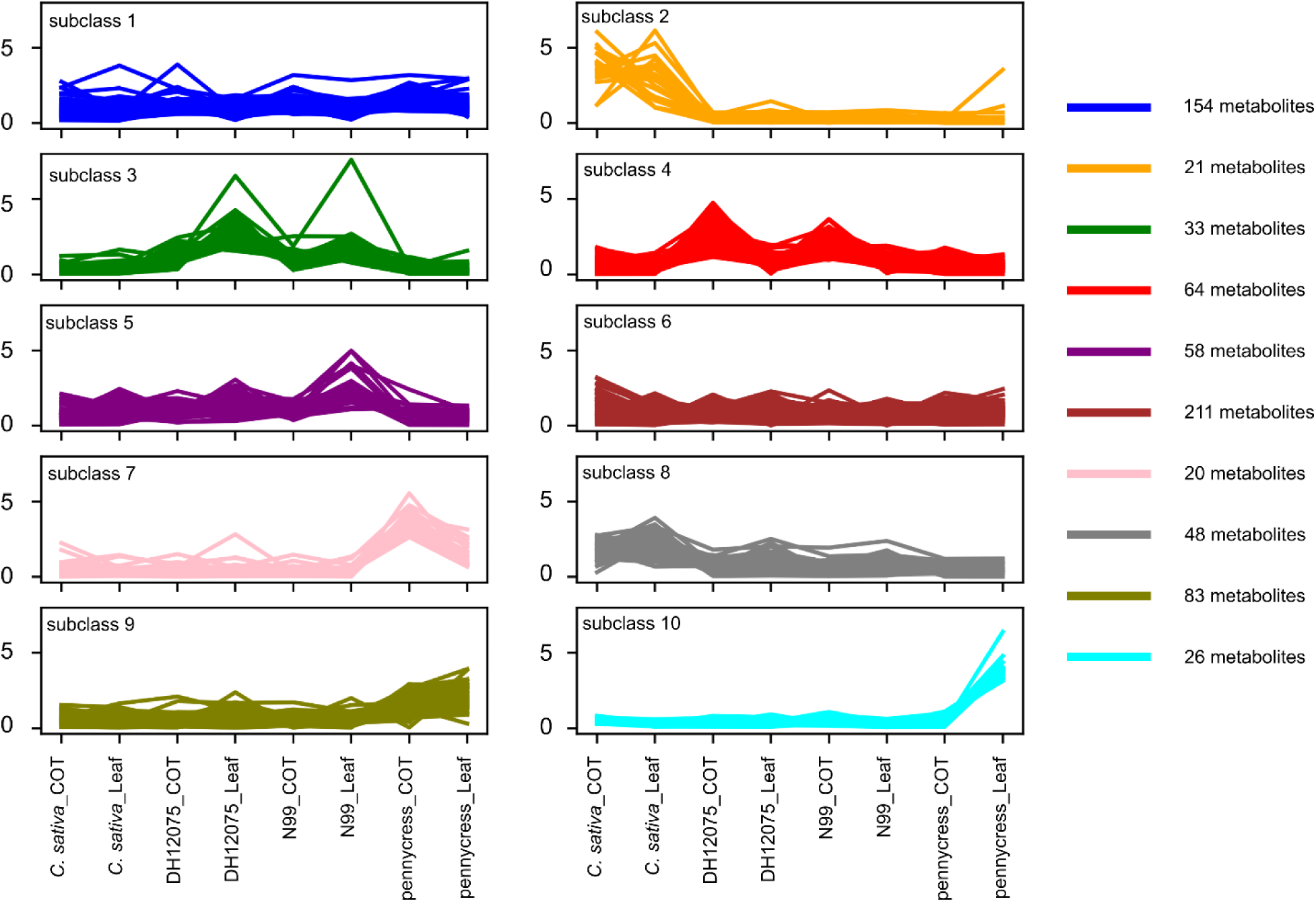
Hierarchical clustering analysis of 718 metabolites based on their relative abundance level separating these metabolites into 10 subclasses. DH12075 and N99 represent two different genotypes of *B. napus*.

**Fig 2.**
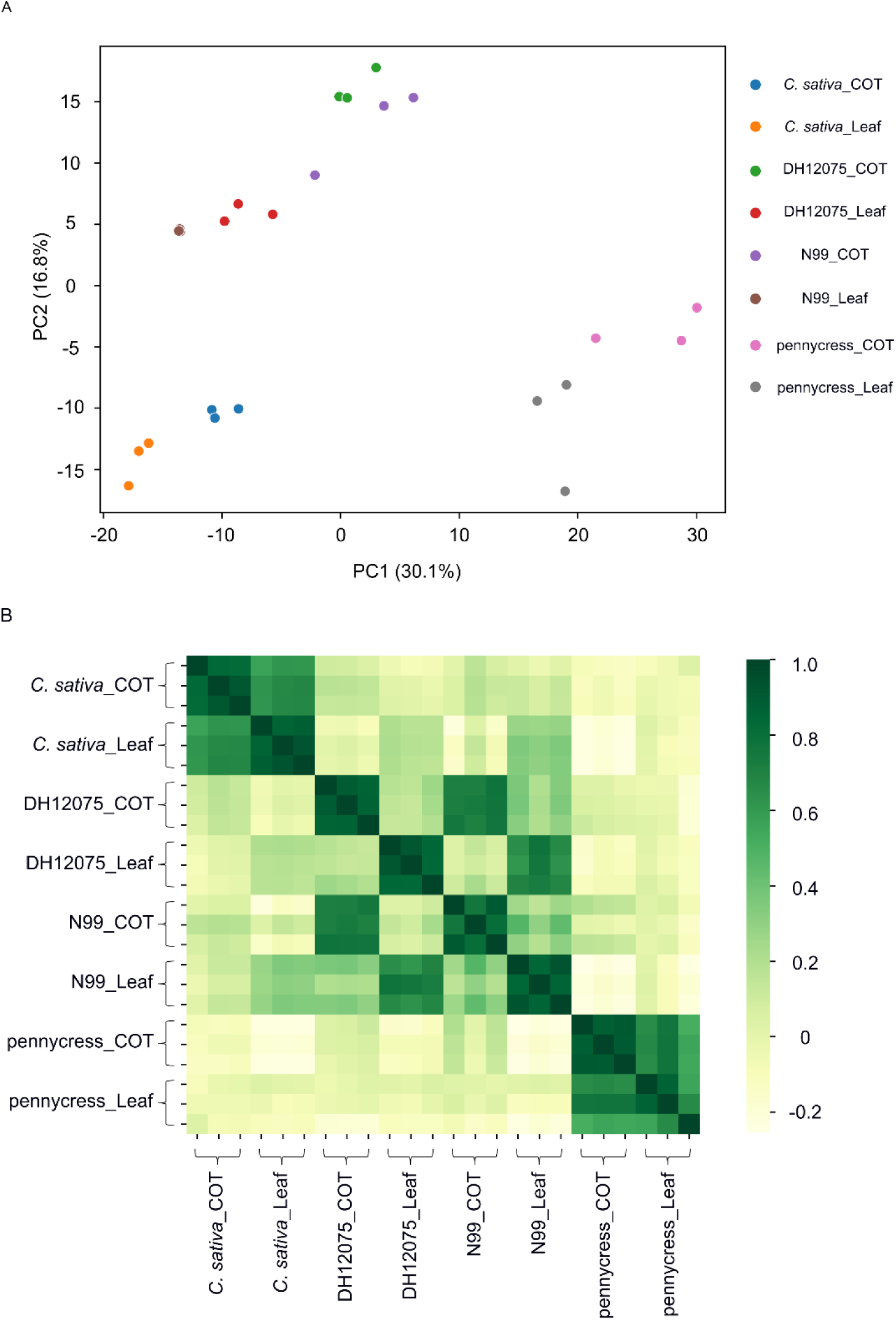
Principal component analysis (PCA) and sample-sample correlation analysis. (A) PCA score plot. (B) Heatmap indicating Pearson correlation coefficients between individual samples.

### 2, PCA analysis and sample-sample correlation

To determine the variability between different groups (species/tissue), as well as the metabolic differences among the three biological replicates for each group, a principal component analysis (PCA) were performed with all detected 718 compounds. Figure 2A showed that the first principal component (PC1) accounted for 30.1% of the variation, and largely separated pennycress from the other two species. Meanwhile the second principal component (PC2) accounted for 16.8% of the variation and separated *B. napus* from *C. sativa* (Figure 2A). The PCA plot shows that for each group, three biological replicates are highly clustered together, suggesting there is a high cohesion within each group. Meanwhile, the samples from the three oilseed crops separated into three distinct areas in the plot, indicating each species possessed a distinct metabolic profile overall. Further, two *B. napus* lines, DH12075 and N99, were closely grouped together, implying there was limited differences between these two genotypes with regards to the metabolic profile (Figure 2A). To further confirm the reproducibility, a sample-sample correlation analysis was performed, where the resultant heatmap showed very high correlation between the three biological replicates for each group. Although, for each species, the metabolic profiles of their cotyledons and leaves were very similar, it was clear from the *B. napus* data that the individual tissues from the two genotypes were more similar to each other than the differences between the genotypes (Figure 2B). Yet, as the main goal was to identify differentially accumulated metabolites at the species-level, results from leaves and cotyledons were combined for each species during subsequent analysis. Additionally, since the two *B. napus* genotypes (i.e. DH12075 and N99) possessed very similar metabolites overall, DH12075 was used to represent the oilseed *B. napus* during the following analysis.

### 3, Pairwise comparisons

To check the overall metabolic differences between these three oilseed crops, we performed pairwise comparisons among them. Each compound was compared between two species by a Student’s t-test, the p-value of which was further corrected through the Benjamini-Hochberg procedure. Significantly different metabolites were selected based on their adjusted p-values with 0.05 as a cut-off value. During the three pairwise comparisons, a total of 267 compounds showed differential abundance when comparing *B. napus* vs. *C. sativa*, 383 for *B. napus* vs. pennycress, and 376 for *C. sativa* vs. pennycress (Figure 3A; Table 2). The comparisons revealed very similar results between *B. napus* vs. pennycress and *C. sativa* vs. pennycress, with the number of their differential metabolites reaching 383 and 376 respectively, among which they shared 268 common metabolites (Table 2; Figure 3B). Meanwhile, the smallest difference was detected in comparisons of *B. napus* vs. *C. sativa*, where only 267 compounds showed differential abundance (170 had higher abundance in *B. napus* while 97 were higher in *C. sativa*) (Figure 3A and B; Table 2).

**Fig 3.**
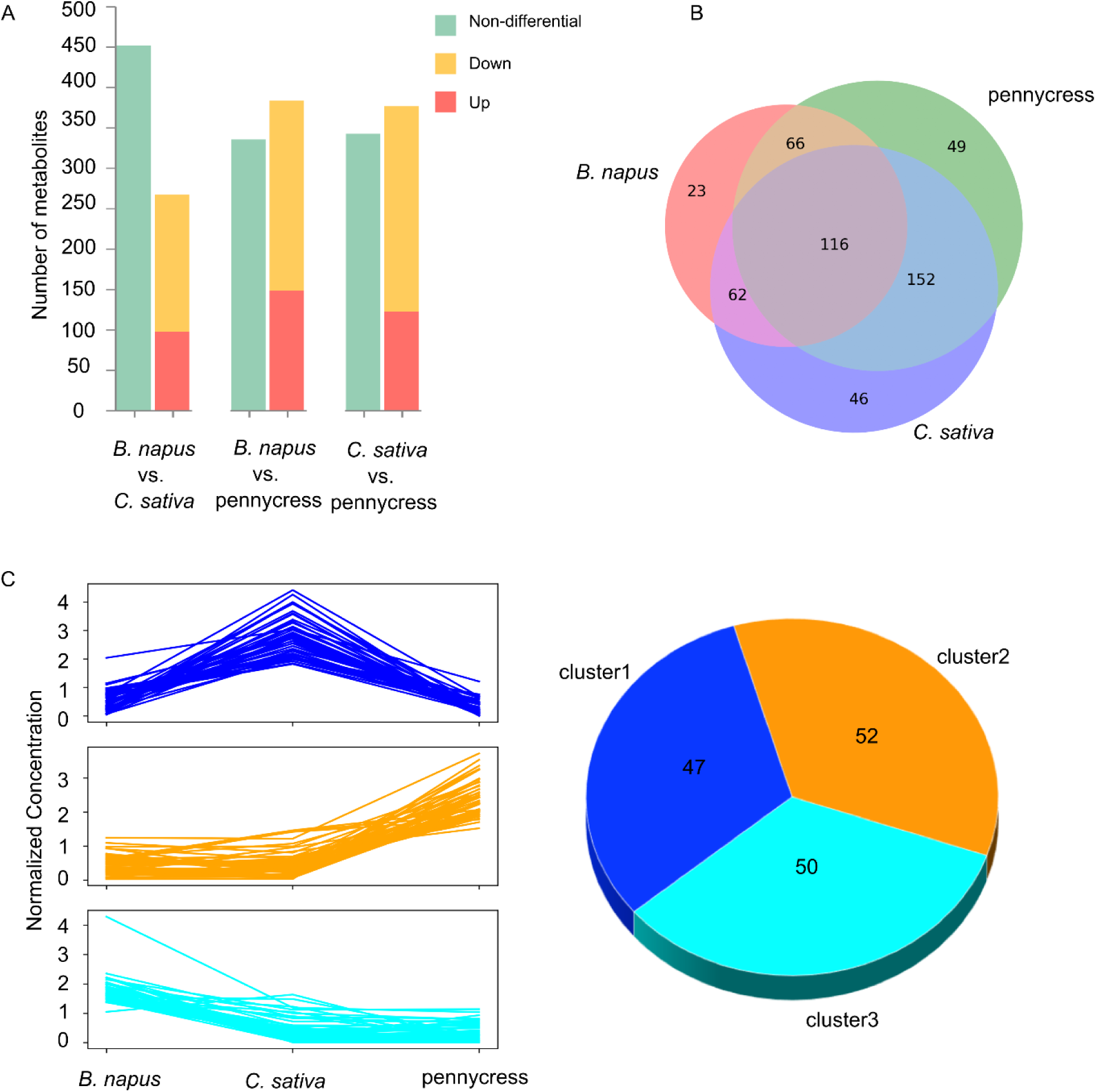
Pairwise comparisons of 718 metabolites between three oilseed crops and clustering of 149 differentially abundant metabolites (DAMs). (A) Bar plots showing results of pairwise species comparisons. (B) Venn diagram showing the overall differential metabolites among three crops. (C) Plot indicating mean value of abundance of the 149 differentially abundant metabolites (DAMs) identified by partial least squares-discriminant analysis (left panel) as well as the number of DAMs belonging to each sub-cluster (right panel).

**Table 2.**
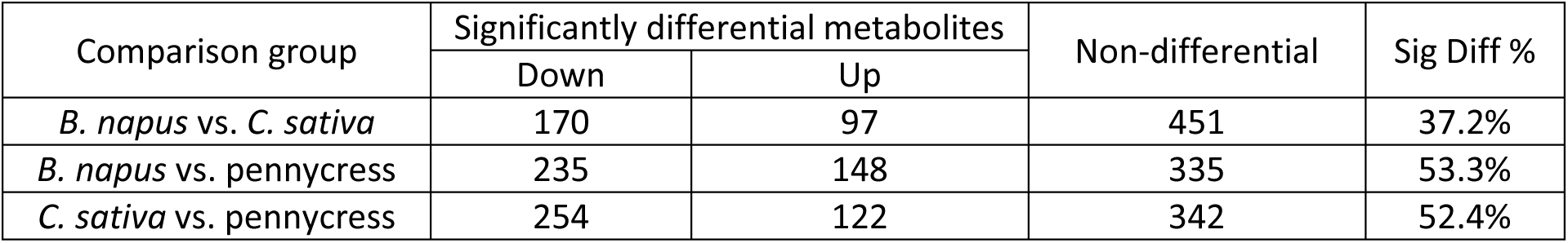
Pairwise comparisons of three oilseed crops.

### 4, PLS-DA identifies differential metabolites

After the explorative analysis above, our main question concerned those metabolites that showed a discriminant pattern among these three oilseed crops. The partial least squares-discriminant analysis (PLS-DA) is an effective method for separating differentially abundant metabolites (DAMs) between different groups in metabolomics data because of its ability to handle highly collinear and noisy data, and maximize the differences between groups (Gromski et al., 2015; Qi et al., 2022). To find the most discriminant metabolites between *B. napus*, *C. sativa* and pennycress, we carried out a PLS-DA with all 718 identified metabolites. According to the variable importance in projection (VIP) scores from PLS-DA, DAMs were initially selected if their VIP scores were higher than 1. Further, all initial DAMs were filtered by an ANOVA to only keep the ones with p-value smaller than 0.05. Thus, DAMs were defined as metabolites with VIP score > 1 and ANOVA p-value < 0.05, which resulted in a total of 149 final DAMs (Supplemental Data2). Based on the relative metabolite abundance, these 149 DAMs were divided into three distinct sub-groups (Figure 3C). Among the three sub-groups, there were distinct metabolite distributions: sub-group 1 comprised 47 metabolites, exhibiting the highest abundance in *C. sativa*; sub-group 2 encompassed 52 metabolites, demonstrating highest concentration in pennycress; 50 metabolites in the 3rd sub-group, displayed the highest concentration in *B. napus* instead (Figure 3C).

To check the top DAMs among these three crops, we selected the 20 metabolites with the highest VIP scores, and plotted these metabolites along with their relative abundance across all three oilseed crops for visualization (Figure 4A). Agmatine, a precursor for polyamine biosynthesis, has the highest VIP score, and its abundance was significantly higher in *C. sativa* compared with the other two oilseed crops (Figure 4A). To further gain knowledge of what biological pathways these 149 compounds were involved in, we performed a functional enrichment analysis according to the Kyoto Encyclopedia of Genes and Genomes (KEGG) database through the pathway enrichment analysis of MetaboAnalyst6 (https://www.metaboanalyst.ca/). As shown in Figure 4B, several amino acid metabolism pathways, including “Tyrosine metabolism”, “Arginine and proline metabolism”, and “Cysteine and methionine metabolism” have very small p-values (p-value < 0.01), which is in line with the large number of overall “Amino Acids & Derivatives” identified in the untargeted metabolomics results (Table 1). Meanwhile, “Isoquinoline alkaloid biosynthesis” also had a small p-value (p-value < 0.05). More noticeably, “Phenylpropanoid biosynthesis” pathway has the smallest p-value and the ensuing “Flavonoid biosynthesis” and “Flavone and flavonol biosynthesis” also have small p-values (p-value = 0.056 and 0.037 respectively).

**Fig 4.**
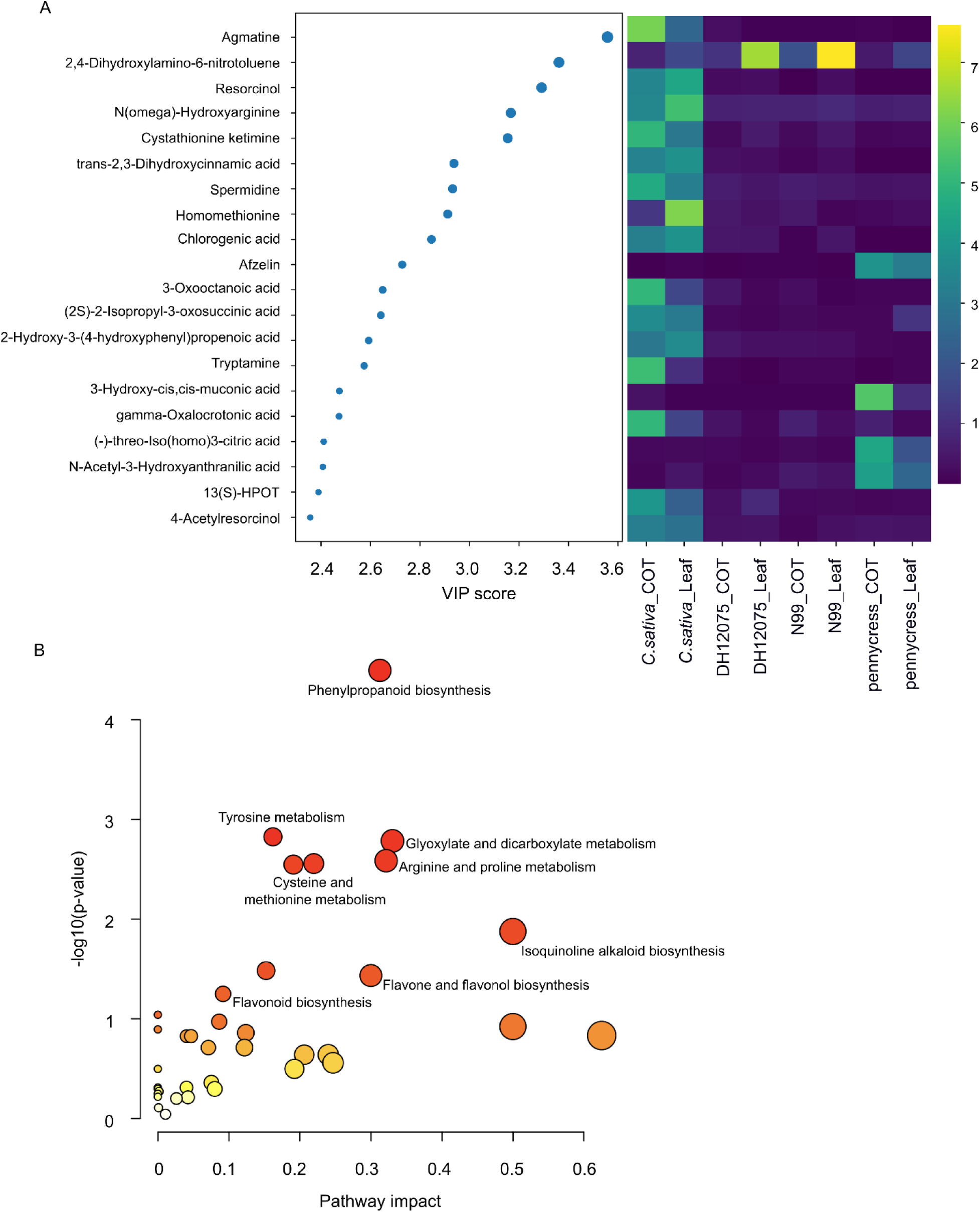
Top twenty differentially abundant metabolites (DAMs) and pathway enrichment analysis. (A) Plot showing 20 metabolites with the highest variable importance in projection (VIP) scores from partial least squares-discriminant analysis (left panel) and their relative abundance across the three oilseed crops (right panel). (B) KEGG pathway enrichment analysis result of 149 differentially abundant metabolites (DAMs).

### 5, Transcriptomic profiles

To investigate the candidate genetic regulators related to the abovementioned DAMs, we performed a concomitant RNA sequencing along with the metabolomics. In order to compare the gene expression between these three oilseed crops, all their genes were first projected onto their corresponding orthologs in *A. thaliana* (see details in Methods). In total, there were 14,025 unique *A. thaliana* genes possessing corresponding orthologous genes in all three oilseed crops, which were selected for further comparison (Supplemental Data3). To identify the differentially expressed genes (DEGs) among these three crop species, a PLS-DA was employed for the 14,025 genes. Under the definition of DEG as “VIP score > 2 and ANOVA p-value < 0.05”, there were 580 DEGs in total. Further, a hierarchical clustering was performed for these 580 DEGs based on their expression profiles. As shown in Figure 5A, these DEGs’ expression patterns varied to a great degree, and the three oilseed crops could be distinguished clearly according to the expression level of these DEGs (Figure 5A). To functionally characterize these identified DEGs, a KEGG enrichment analysis was performed, where pathways “Glutathione metabolism”, “2-Oxocarboxylic acid metabolism”, and “Cysteine and methionine metabolism” etc. were significantly enriched (Figure 5B).

**Fig 5.**
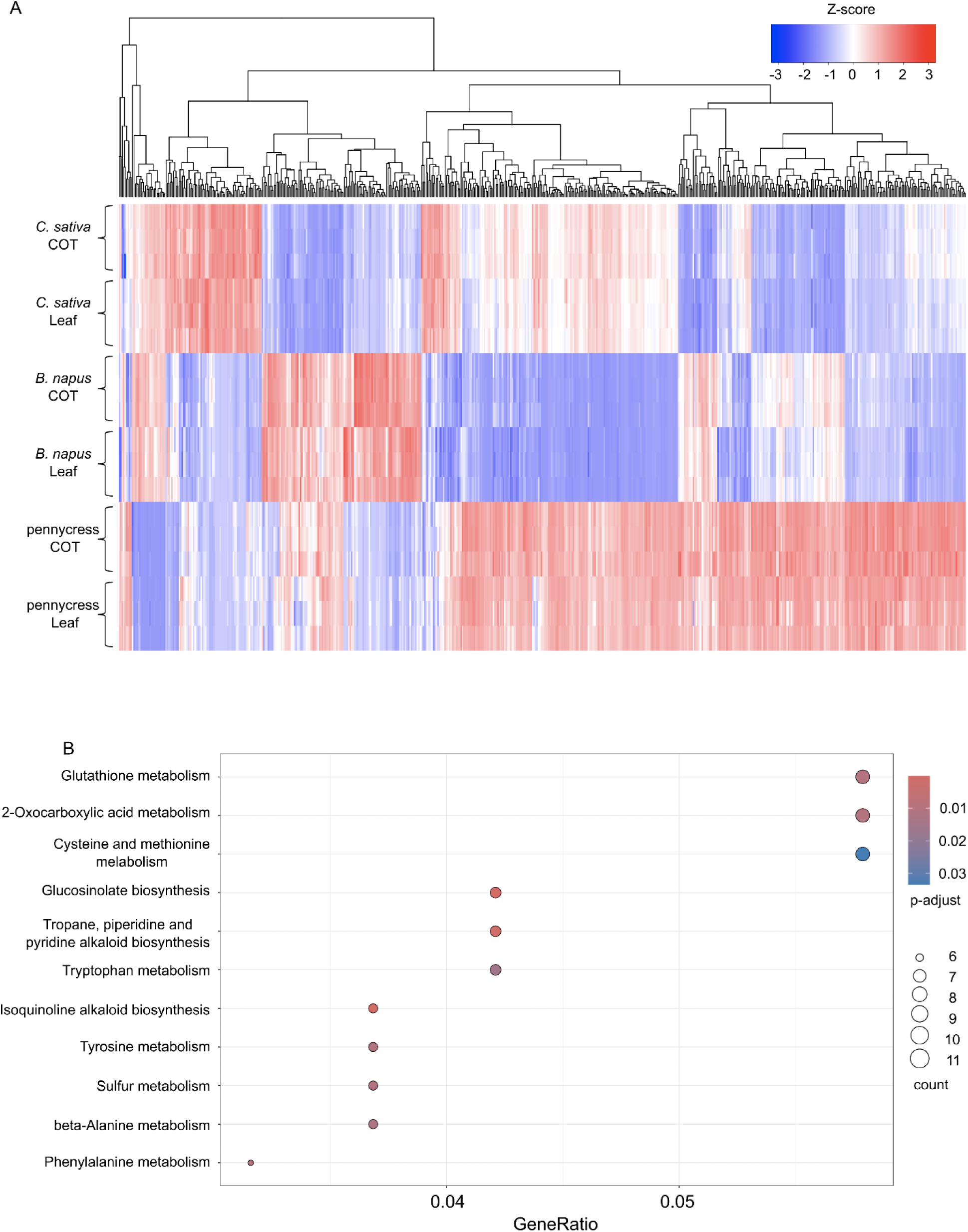
Differentially expressed genes (DEGs) and KEGG pathway enrichment analysis. (A) Heatmap showing the hierarchical clustering analysis of all 580 differentially expressed genes (DEGs). (B) Dot plot showing the KEGG pathway enrichment analysis result. Color bar indicating the p-values from Fisher’s exact test, and dots size indicating the number of genes belonging to the each pathway.

### 6, Integration of genes and metabolites in phenylpropanoid pathway

As the 149 DAMs above were mainly attributed to the “Phenylpropanoid biosynthesis” pathway (Figure 4B), to further examine this pathway in more detail, we selected all DAMs that were classified as phenylpropanoids or flavonoids by the KEGG database, and compared their relative abundances across these three oilseed crops. The compounds were involved in general phenylpropanoid pathway (e.g. *p*-Coumaric acid), flavonoid pathway (e.g. Afzelin, Vitexin, Kaempferol-3-O-galactoside), and phenolic acid pathway (e.g. Chlorogenic acid). The relative abundances of these phenylpropanoids varied to a great extent, for example, *C. sativa* contains the highest levels of Chlorogenic acid and cis-3,4-Leucopelargonidin, while *B. napus* contains the highest levels of Kaempferol-3-O-galactoside, trans-Ferulic acid, and *p*-Coumaroyl quinic acid, meanwhile Vitexin and Afzelin accumulated the most in pennycress (Figure 6).

**Fig 6.**
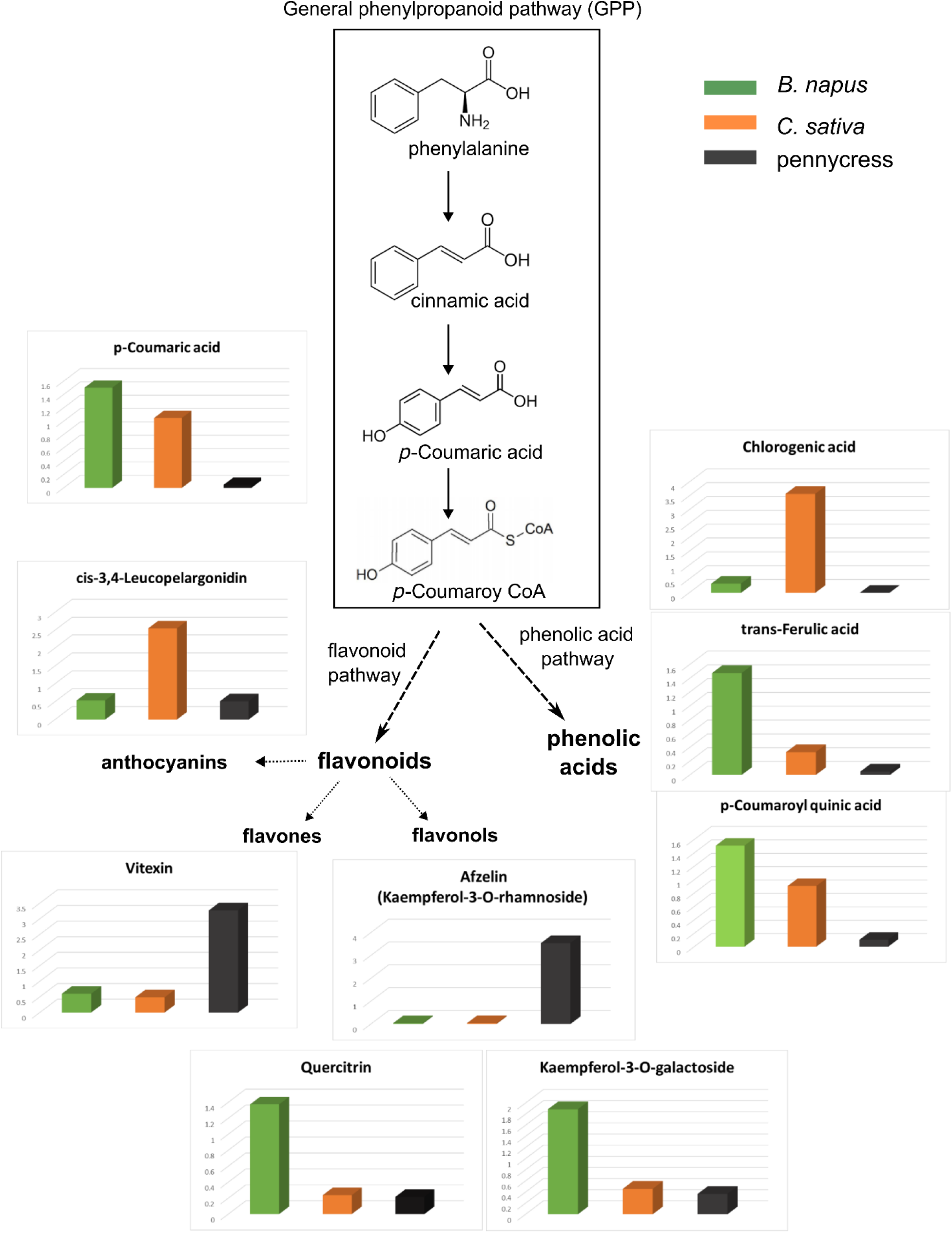
Phenylpropanoids, flavonoids, and phenolic acids in three oilseed crops. 3D boxplots showing relative abundance of individual metabolite. *B. napus* was labelled in green, *C. sativa* in orange and pennycress in grey.

To identify genes that were responsible for the abovementioned phenylpropanoids, multiple putative regulatory genes involved in phenylpropanoid biosynthesis were selected to compare their expression levels. As might be expected, there was great variation in expression levels for phenylpropanoid biosynthetic genes between these three oilseed crops (Figure 7); for example, homologs of flavonol 3’-hydroxylase (F3’H/TT7) and dihydroflavonol reductase (DFR/TT3) were predominantly expressed in *C. sativa*, meanwhile homologs of leucoanthocyanidin dioxygenase (LDOX/ANS/TT18) were mainly expressed in *B. napus*, these expression differences were in line with the high accumulation of cis-3,4-Leucopelargonidin in *C. sativa* (Figure 6 and Figure 7). However, given the limitation of untargeted metabolomics, there were only a small amount of identified phenylpropanoids, which makes it difficult to link the variation in gene expression directly to the corresponding change in phenylpropanoids.

**Fig 7.**
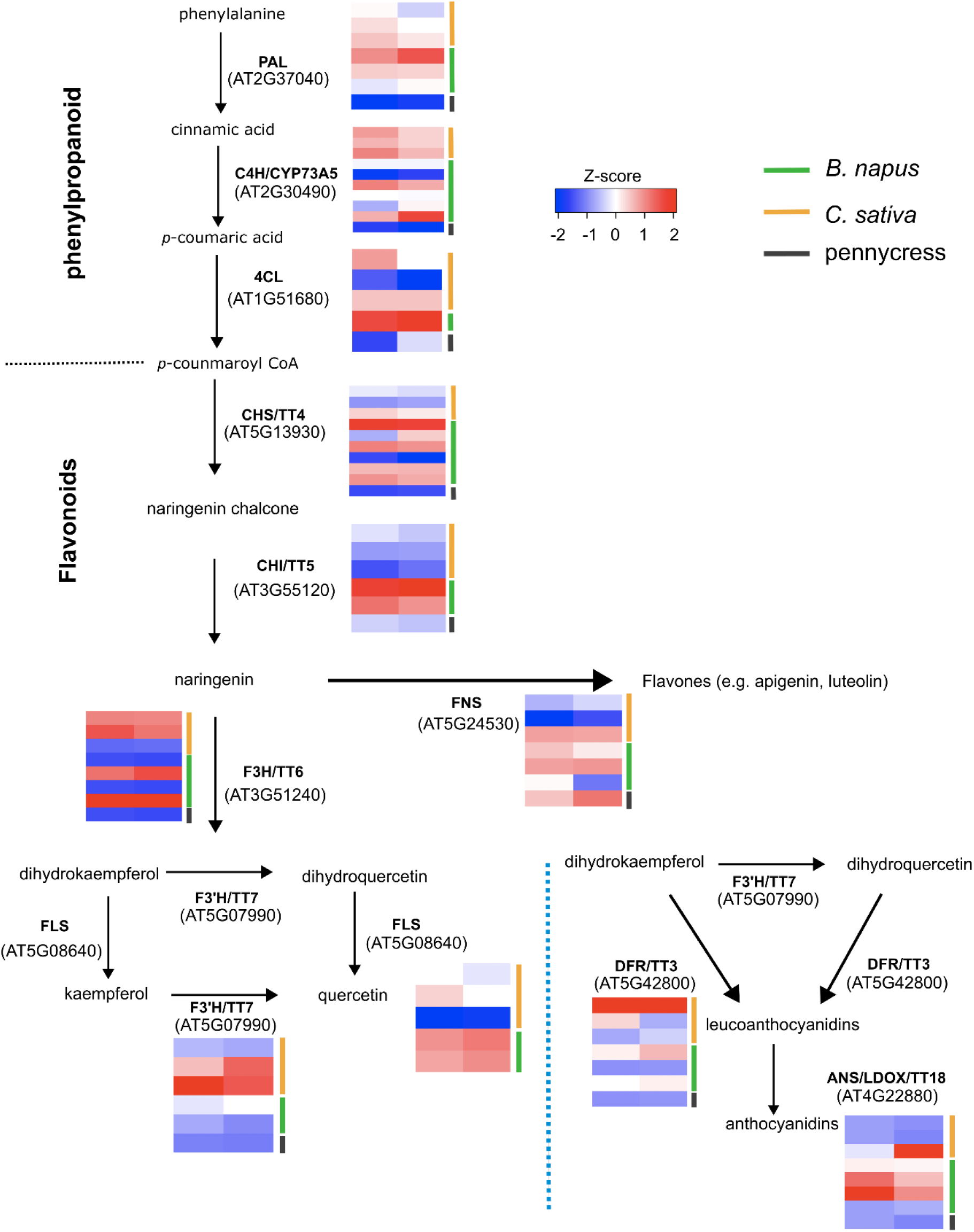
Expression of the genes involved in pathway of phenylpropanoid and flavonoid biosynthesis. The genes’ standard scores (Z-scores) were used for comparison and expression levels are illustrated with white, blue or red blocks. Left column corresponds to expression in cotyledon tissue, and right column corresponds to expression in leaf tissue. Orthologous genes from *B. napus*, *C. sativa* and pennycress were labelled in green, orange and grey respectively. The dotted blue line separates the alternative anthocyanin biosynthesis pathway. PAL, phenylalanine ammonia-lyase; C4H/CYP73A5, cinnamic acid 4-hydroxylase; 4CL, 4-coumarate-CoA ligase; CHS/TT4, chalcone synthase; CHI/TT5, chalcone isomerase; FNS, Flavone synthase; F3H/TT6, flavonol 3-hydroxylase; F30H/TT7, flavonol 30-hydroxylase; FLS, flavonol synthase; DFR/TT3, dihydroflavonol reductase; ANS/LDOX/TT18, leucoanthocyanidin dioxygenase.

To explore the 580 DEGs in a more general manner, co-expressed gene modules were identified through weighted gene co-expression network analysis (WGCNA) (Langfelder & Horvath, 2008). As shown in Figure 8A, these 580 DEGs were divided into 7 separate modules based on their expression levels (Supplemental Data4). According to the eigengene expression of each module, module yellow, green and blue were *B. napus*, *C. sativa*, and pennycress specific, respectively (Figure 8B). Interestingly, *C. sativa*-specific module green was highly correlated with the 47 DAMs in sub-group 1, which accumulated the highest abundance in *C. sativa* (Figure 3C; Figure 8C). Similarly, *B. napus*-specific module yellow and pennycress-specific module blue were strongly correlated with 50 DAMs in sub-group 2 and 52 DAMs in sub-group 3 (Figure 3C; Figure 8C), thus suggesting that the species-specific gene expression identified among these three oilseed crops might contribute to their distinct metabolic profiles. To mine potential candidate genes associated with variation of the abovementioned phenylpropanoids, we further examined the correlations between the identified phenylpropanoids and the 580 DEGs. Pearson’s correlation coefficients showed that 157 DEGs were significantly related to 9 phenylpropanoids (R^2^ > 0.9), whereat 108 pairs were positively correlated, and 56 pairs were negatively correlated (Supplemental Data5), these 157 DEGs were highly enriched in GO terms “alpha-amino acid metabolic process” and “secondary metabolite biosynthesis process”.

**Fig 8.**
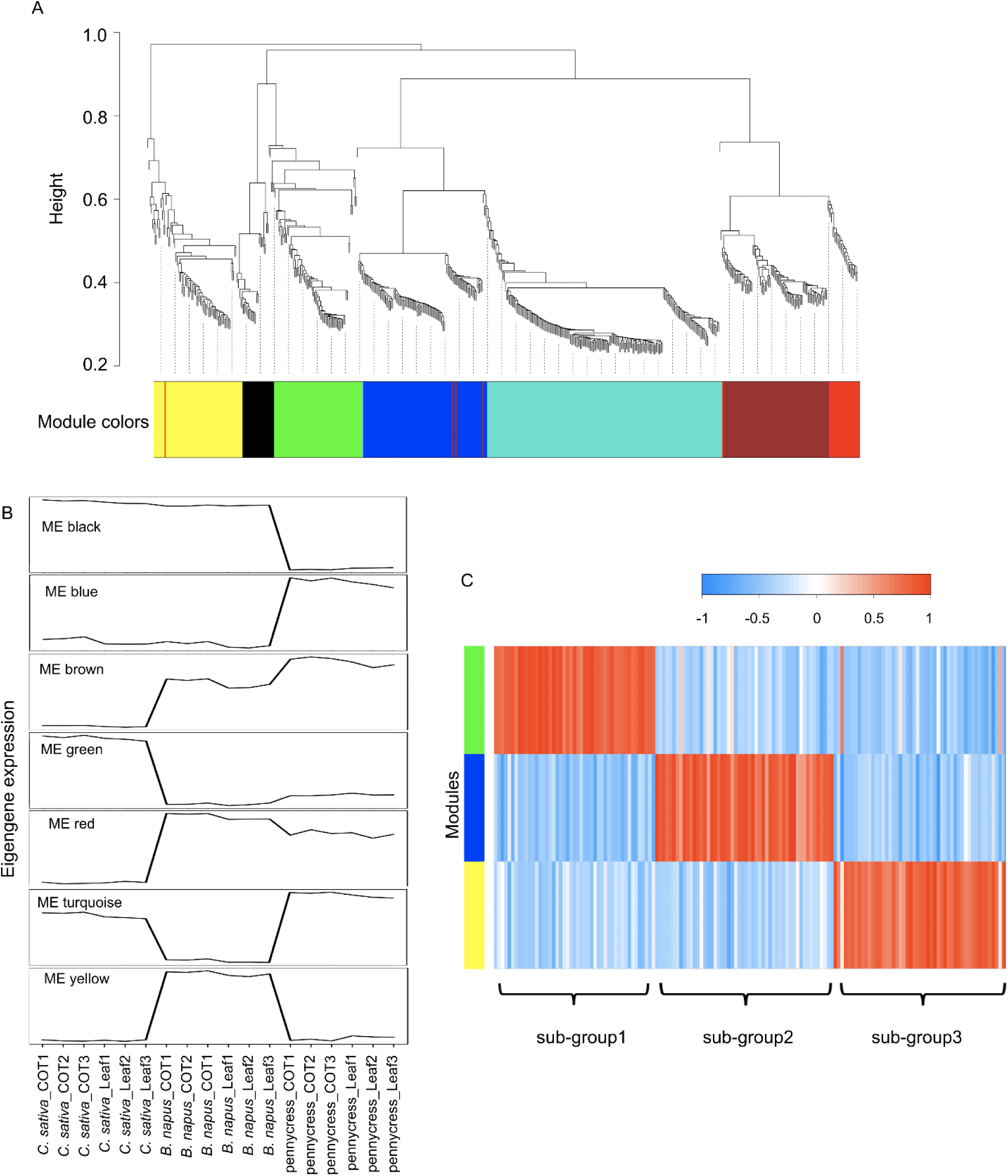
WGCNA identifies co-expressed gene modules. (A) dendrogram indicating 7 distinct modules and different colors were assigned to each individual module. (B) Plot showing eigengene expression of each module across the three species (C) heatmap showing the correlation of 149 DAMs from Figure 3C with co-expression module green, blue and yellow.

In summary, to document the metabolic profiles of *B. napus*, *C. sativa*, and field pennycress, a total of 718 metabolites including flavonoids, phenylpropanoids, amino acids and derivatives, were characterized through untargeted metabolomics analysis. Moreover, 149 DAMs were identified through PLS-DA, indicating the different metabolic profiles of these three crops were mainly attributed to phenylpropanoids. Further integration with transcriptomics revealed a great variation of the candidate genes’ expression between these three species. Our data provides a novel resource for understanding the metabolism of these three oilseed crops at the molecular level.

## Discussion

### Untargeted metabolomics analyses reveals large metabolic variation of three oilseed crops in unstressed leaf tissue

Although, metabolomics has been widely used to study the metabolites of plants (Allwood et al., 2021; Wang et al., 2016; Yan et al., 2022), in oilseed crops, most attention has focused on dissecting the metabolic profile of their seeds, in order to further improve the oil quality. For example, Boutet et al. (2022) revealed a large diversity of specialized metabolites in Camelina seeds through untargeted metabolomics and lipidomics (Boutet et al., 2022). Li et al. (2023) utilized targeted metabolomics to study the oil content of *Brassica napus* seeds, which identified marker metabolites that were correlated with oil content (Li et al., 2023). However, there is very limited knowledge about the metabolites in the leaf and/or cotyledon tissues of these oilseed crops. Metabolites play key roles in plants’ physiological and biochemical responses to environmental stimuli, and the accumulation of metabolites also has been shown to vary to a great extent in different species (Qi et al., 2021; Wang et al., 2016). Understanding the metabolite differences between species in tissues such as cotyledons and young leaves, which are often the first to recognize new threats, could provide insights into why some species are more resilient than others. In this instance, two novel oilseed crops *C. sativa* and *T. arvense* that have often been associated with higher tolerance to a number of biotic and abiotic stresses compared to their more widely grown relative *B. napus* were used to test this hypothesis.

In this study, we detected more than 3,465 metabolic peak signals through LC-MS-based untargeted metabolomics (Supplemental Data6). Underlying the current limitations of these technologies, only 718 metabolites could be identified given that the majority of metabolites in the plant kingdom are still unknown or unannotated (Wang et al., 2019; Yonekura-Sakakibara & Saito, 2009). The analysis of these 718 identified metabolites revealed great variation between these three oilseed crops, indicating a species-specific accumulation pattern of metabolites (Figure 1), which was further supported by the PCA plot that showed complete separation of these three species (Figure 2A). The 149 DAMs identified by PLS-DA were highly enriched in “Phenylpropanoid biosynthesis”, and amino acid metabolism pathways (Figure 4B), suggesting that both phenylpropanoids and amino acids play important roles in distinguishing these three oilseed crops.

### Phenylpropanoids and flavonoids in the Brassicaceae family

Flavonoids are a subgroup of phenylpropanoids, which are an important class of plant secondary metabolites synthesized via the shikimate pathway from phenylalanine (Dixon et al., 2002; Fraser & Chapple, 2011; Vogt, 2010). In addition to flavonoids, phenylpropanoids contain other different subgroups, including lignins, phenolic acids, stilbenes, and coumarins (Deng & Lu, 2017; Vogt, 2010). Both phenylpropanoids and flavonoids have been shown to play important roles in plant defense against abiotic and/or biotic stresses (Ramaroson et al., 2022; Sharma et al., 2019). Our current results indicated that there was great variation in the phenylpropanoids, including flavonoids and phenolic acids, between these three oilseed crops, e.g. *C. sativa* contains high amount of chlorogenic acid (Figure 4B and 6). However, given the limitation of using untargeted metabolomics, where a significant portion of the detected peaks are not characterized, only a small number of phenylpropanoids were identified, which hinders our understanding of where the different metabolic pathways diverge in the three species. A previous study by Onyilagha *et al*. (2003) using a targeted approach, presented a more detailed view of leaf flavonoid distribution among different crops from the Brassicaceae family through Thin-layer Chromatography (TLC) and high-performance liquid chromatography (HPLC), whereat quercetin was the only type of flavonoid detected in *C. sativa*; while *B. napus* accumulated at least two types of flavonols, with kaempferol as the major type and quercetin in a small concentration (Onyilagha et al., 2003). A further study confirmed that *B. napus* leaves mainly contained kaempferols, while *C. sativa* accumulated large amount of quercetins instead (Onyilagha et al., 2012). Interestingly, high kaempferols in *B. napus* and high quercetins in *C. sativa* is in line with our transcriptomic data, where the homologs of TT7, which encodes a flavonol 3’-hydroxylase to convert kaempferols to quercetins, were predominately expressed in *C. sativa*, (Figure 7). Genetic editing and kinetic analysis of TT7 homologs in *C. sativa* could be useful for further verification of their functions.

### High polyamines endow *C. sativa* with resistance against broad range of abiotic stresses?

Compared with other Brassicaceae oilseed species, *C. sativa* has been shown to tolerate a broad range of abiotic stresses, including but not limited to drought, freezing, lodging, and salinity (Berti et al., 2016; Čanak et al., 2020; Gao et al., 2018; Matthees et al., 2018). However, the mechanisms underlying these tolerances of *C. sativa* are largely unknown. Our results here indicated that there were particularly high content of polyamines in *C. sativa* compared to other two crops. Noticeably there are two polyamines in the top 20 DAMs, agmatine and spermidine, both of which accumulate in *C. sativa* (Figure 4A). Interestingly, both agmatine and spermidine are involved in the “Arginine and proline metabolism” pathway (Figure 4B), where agmatine is an intermediate metabolite formed by arginine decarboxylase (ADC) from arginine, further agmatine is used to generate spermidine through sequential reactions (Chen et al., 2018).

More importantly, both agmatine and spermidine have been shown to play important roles during plant defense against a variety of abiotic stresses, including drought, freezing and salinity etc. (Gupta et al., 2013; Kasukabe et al., 2004; Seifi & Shelp, 2019; Shao et al., 2022). For example, Kasukabe et al., 2004 showed that overexpression of spermidine synthase in *A. thaliana* significantly increased the spermidine content in leaves, and provided enhanced tolerance to various abiotic stresses (Kasukabe et al., 2004). Therefore, it’s reasonable to speculate that the high content of polyamines (e.g. agmatine and spermidine) in *C. sativa* is likely to contribute to its resistance to a broad range of abiotic stresses. In the future, quantifying and comparing other major polyamines (e.g. diamine putrescine and tetraamine spermine) between these oilseed crops, as well as genetic modification of related polyamine biosynthetic genes in *C. sativa*; for example, targeted knock-out of agmatine and spermidine biosynthetic genes, encoding arginine decarboxylase (ADC) and Spd synthase respectively, woud help us better elucidate the function of polyamines in the broad abiotic resistances of *C. sativa*.

## Materials and Methods

### Plant Materials

*Brassica napus* (DH12075 and N99), *Camelina sativa* (DH55), and pennycress (line collected from field) were grown in a growth chamber (22°, 16 h light /18°, 8 h dark cycles) for up to 2 weeks after germination. Three biological replicates of cotyledons and first true leaves for each species were collected and ground into powder in liquid nitrogen, then stored at -80° freezer for metabolomic and transcriptomic analyses.

### Metabolite extraction and LC-MS analysis

Untargeted metabolomic analyses was carried out by the Metabolomics Innovation Centre (University of Alberta, CA). In short, for each sample, 40 mg tissue powder was used to extract metabolites, where 6 ceramic beads were placed in the sample vials, and 500 µL LC-MS grade MeOH/water (4:1 v/v) was added before homogenizing at 4.5 m/s for 15 seconds. Then, the homogenates were incubated at -20°C for 10 minutes and centrifuged at 15,000 g for 10 minutes, after which the supernatants were carefully transferred into new vials and completely dried. Sample extracts were then re-suspended in 30 µL LC-MS grade water before chemical isotope labeling. The subsequent LC-MS analyses were carried out with a Thermo Scientific Vanquish LC linked to Bruker Impact II QTOF Mass Spectrometer (Bruker, Germany) using the eclipse plus reversed-phase C18 column (150 x 2.1 mm,1.8 µm particle size; Agilent) at 40°C with a flow rate of 400 μL/min.

### Metabolite Quantification and Data Analysis

The raw LC-MS data were first processed using DataAnalysis 4.4 (Bruker), which then exported data to IsoMS Pro 1.2.16 (Nova Medical Testing Inc, CA) for quality check and processing with the following parameters: minimum m/z: 220, maximum m/z: 1000, saturation intensity: 20,000,000, retention time tolerance: 9s, mass tolerance: 10 ppm. Further, data files were filtered out peak pairs present in less than 80% of samples, and normalized by ratio of total useful signal. Metabolite Identification were carried out against the NovaMT Metabolite Database v3.0 (The Metabolomics Innovation Centre, CA).

The hierarchical clustering analysis, principle component analysis (PCA), and partial least squares-discriminant analysis (PLS-DA) were carried out through scikit-learn (version 1.2.2) in Python (version 3.10.9). KEGG pathway functional enrichment analysis was performed via MetaboAnalyst6 (https://www.metaboanalyst.ca/) using the *A. thaliana* database.

### RNA sequencing and data analysis

Total RNA of each sample was extracted using RNAeasy plant mini Kit (Qiagen) according to the manufacturer’s instructions. The quality of total RNA were examined by BioAnalyzer with RNA 6000 Nano Kit (Agilent) to ensure RNA integrity number value > 7. Total RNA were used to prepare cDNA libraries following Illumina Stranded mRNA Prep guide, further 150 bp paired-end sequencing was performed on the NovaSeq 6000 platform (Illumina).

The raw RNA-seq data were first filtered using Trimmomatic (version 0.32) to remove adapter and low-quality sequences. Then clean reads were aligned to the corresponding reference transcriptomes of *B. napus* (DH12075 v3.1; cruciferseq.ca), *C. sativa* (Kagale et al., 2014) and pennycress (Nunn et al., 2022) respectively using Salmon (version 1.10.0). Subsequently, tximport (version 1.30.0) was used to obtain the gene expression levels as TPM (transcripts per million), which were further normalized through log2 transformation.

Syntelog tables of the *A. thaliana* orthologous genes were collected for *B. napus* (DH12075 v3.1; cruciferseq.ca) and *C. sativa* (Kagale et al., 2014). Homologous pairs between pennycress and *A. thaliana* were obtained through Reciprocal BLAST Hits (RBH). To make the expression comparable across three different species, all genes were projected to their corresponding *A. thaliana* orthologs for each species. Given the polyploidy of *B. napus* and *C. sativa*, they contain multiple copies of orthologs for each *A. thaliana* gene. To simplify the comparison, for each *A. thaliana* gene, the highest expressed orthologous gene in *B. napus* and *C. sativa* was selected as the representative gene for subsequent comparison.

To identify the differentially expressed genes (DEGs) between these three species, a partial least squares discriminant analysis (PLS-DA) was performed through scikit-learn (version 1.2.2) in Python (version 3.10.9). One-way ANOVA was performed to calculate the raw p-values, which were further corrected by the false discovery rate (FDR) method for the multiple comparisons. The KEGG enrichment analysis of DEGs was performed using clusterProfiler (version 4.11.0) in R (version 4.3.3). The 580 DEGs were used to construct a co-expression network using the WGCNA package in R with the blockwiseModules function and parameters “power= 12, maxBlockSize = 5000, networkType = "signed", TOMType = "signed", minModuleSize = 10, mergeCutHeight = 0.15” to construct a signed network and generate co-expressed gene modules (Langfelder & Horvath, 2008).

### Integration of metabolome and transcriptome

For integration of metabolomic and transcriptomic data, Pearson correlation coefficients were calculated for each gene-metabolite pair using the ‘cor’ package in R (version 4.3.3). The gene-metabolite network was constructed with R2 > 0.9, where the nodes correspond to genes/metabolites, and the edges between the nodes represent the correlation coefficients calculated (Supplemental Data5).

## DATA AVAILABILITY STATEMENT

The raw RNA-seq read data were deposited to the Gene Expression Omnibus (GEO) with accession number GSE279986.

## ACKNOWLEDGEMENTS

This work was supported by Saskatchewan Agricultural Development Fund (ADF).

## References

Allwood, J. W., Williams, A., Uthe, H., van Dam, N. M., Mur, L. A. J., Grant, M. R., & Petriacq, P. (2021). Unravelling Plant Responses to Stress-The Importance of Targeted and Untargeted Metabolomics. Metabolites, 11(8). 10.3390/metabo11080558

Arbona, V., Manzi, M., Ollas, C., & Gomez-Cadenas, A. (2013). Metabolomics as a tool to investigate abiotic stress tolerance in plants. Int J Mol Sci, 14(3), 4885–4911. 10.3390/ijms14034885

Arias, C. L., Garcia Navarrete, L. T., Mukundi, E., Swanson, T., Yang, F., Hernandez, J., Grotewold, E., & Alonso, A. P. (2023). Metabolic and transcriptomic study of pennycress natural variation identifies targets for oil improvement. Plant Biotechnol J, 21(9), 1887–1903. 10.1111/pbi.14101

Bartwal, A., Mall, R., Lohani, P., Guru, S. K., & Arora, S. (2013). Role of Secondary Metabolites and Brassinosteroids in Plant Defense Against Environmental Stresses. Journal of Plant Growth Regulation, 32(1), 216–232. 10.1007/s00344-012-9272-x

Berti, M., Gesch, R., Eynck, C., Anderson, J., & Cermak, S. (2016). Camelina uses, genetics, genomics, production, and management. Industrial Crops and Products, 94, 690–710. 10.1016/j.indcrop.2016.09.034

Boutet, S., Barreda, L., Perreau, F., Totozafy, J. C., Mauve, C., Gakiere, B., Delannoy, E., Martin-Magniette, M. L., Monti, A., Lepiniec, L., Zanetti, F., & Corso, M. (2022). Untargeted metabolomic analyses reveal the diversity and plasticity of the specialized metabolome in seeds of different Camelina sativa genotypes. Plant J, 110(1), 147–165. 10.1111/tpj.15662

Čanak, P., Jeromela, A. M., Vujošević, B., Kiprovski, B., Mitrović, B., Alberghini, B., Facciolla, E., Monti, A., & Zanetti, F. (2020). Is Drought Stress Tolerance Affected by Biotypes and Seed Size in the Emerging Oilseed Crop Camelina? Agronomy, 10(12), 1856. https://www.mdpi.com/2073-4395/10/12/1856

Castro-Moretti, F. R., Gentzel, I. N., Mackey, D., & Alonso, A. P. (2020). Metabolomics as an Emerging Tool for the Study of Plant-Pathogen Interactions. Metabolites, 10(2). 10.3390/metabo10020052

Chen, D., Shao, Q., Yin, L., Younis, A., & Zheng, B. (2018). Polyamine Function in Plants: Metabolism, Regulation on Development, and Roles in Abiotic Stress Responses. Front Plant Sci, 9, 1945. 10.3389/fpls.2018.01945

Chhajed, S., Mostafa, I., He, Y., Abou-Hashem, M., El-Domiaty, M., & Chen, S. (2020). Glucosinolate Biosynthesis and the Glucosinolate–Myrosinase System in Plant Defense. Agronomy, 10(11). 10.3390/agronomy10111786

Czerniawski, P., Piasecka, A., & Bednarek, P. (2021). Evolutionary changes in the glucosinolate biosynthetic capacity in species representing Capsella, Camelina and Neslia genera. Phytochemistry, 181, 112571. 10.1016/j.phytochem.2020.112571

Del Carmen Martinez-Ballesta, M., Moreno, D. A., & Carvajal, M. (2013). The physiological importance of glucosinolates on plant response to abiotic stress in Brassica. Int J Mol Sci, 14(6), 11607–11625. 10.3390/ijms140611607

Deng, Y., & Lu, S. (2017). Biosynthesis and Regulation of Phenylpropanoids in Plants. Critical Reviews in Plant Sciences, 36(4), 257–290. 10.1080/07352689.2017.1402852

Dixon, R. A., Achnine, L., Kota, P., Liu, C. J., Reddy, M. S., & Wang, L. (2002). The phenylpropanoid pathway and plant defence-a genomics perspective. Mol Plant Pathol, 3(5), 371–390. 10.1046/j.1364-3703.2002.00131.x

Erb, M., & Kliebenstein, D. J. (2020). Plant Secondary Metabolites as Defenses, Regulators, and Primary Metabolites: The Blurred Functional Trichotomy. Plant Physiol, 184(1), 39–52. 10.1104/pp.20.00433

Fraser, C. M., & Chapple, C. (2011). The phenylpropanoid pathway in Arabidopsis. Arabidopsis Book, 9, e0152. 10.1199/tab.0152

Gao, L., Caldwell, C. D., & Jiang, Y. (2018). Photosynthesis and Growth of Camelina and Canola in Response to Water Deficit and Applied Nitrogen. Crop Science, 58(1), 393–401. 10.2135/cropsci2017.07.0406

Gromski, P. S., Muhamadali, H., Ellis, D. I., Xu, Y., Correa, E., Turner, M. L., & Goodacre, R. (2015). A tutorial review: Metabolomics and partial least squares-discriminant analysis--a marriage of convenience or a shotgun wedding. Anal Chim Acta, 879, 10–23. 10.1016/j.aca.2015.02.012

Gugel, R. K., & Falk, K. C. (2006). Agronomic and seed quality evaluation of Camelina sativa in western Canada. Canadian Journal of Plant Science, 86(4), 1047–1058. 10.4141/p04-081

Gupta, K., Dey, A., & Gupta, B. (2013). Plant polyamines in abiotic stress responses. Acta Physiologiae Plantarum, 35(7), 2015–2036. 10.1007/s11738-013-1239-4

Halkier, B. A., & Gershenzon, J. (2006). Biology and biochemistry of glucosinolates. Annu Rev Plant Biol, 57, 303–333. 10.1146/annurev.arplant.57.032905.105228

Hanada, K., Zou, C., Lehti-Shiu, M. D., Shinozaki, K., & Shiu, S. H. (2008). Importance of lineage-specific expansion of plant tandem duplicates in the adaptive response to environmental stimuli. Plant Physiol, 148(2), 993–1003. 10.1104/pp.108.122457

He, M., He, C. Q., & Ding, N. Z. (2018). Abiotic Stresses: General Defenses of Land Plants and Chances for Engineering Multistress Tolerance. Front Plant Sci, 9, 1771. 10.3389/fpls.2018.01771

Kagale, S., Koh, C., Nixon, J., Bollina, V., Clarke, W. E., Tuteja, R., Spillane, C., Robinson, S. J., Links, M. G., Clarke, C., Higgins, E. E., Huebert, T., Sharpe, A. G., & Parkin, I. A. (2014). The emerging biofuel crop Camelina sativa retains a highly undifferentiated hexaploid genome structure. Nat Commun, 5, 3706. 10.1038/ncomms4706

Kasukabe, Y., He, L., Nada, K., Misawa, S., Ihara, I., & Tachibana, S. (2004). Overexpression of Spermidine Synthase Enhances Tolerance to Multiple Environmental Stresses and Up-Regulates the Expression of Various Stress-Regulated Genes in Transgenic Arabidopsis thaliana. Plant and Cell Physiology, 45(6), 712–722. 10.1093/pcp/pch083

Kirkegaard, J. A., Lilley, J. M., Berry, P. M., & Rondanini, D. P. (2021). Chapter 17 - Canola. In V. O. Sadras & D. F. Calderini (Eds.), Crop Physiology Case Histories for Major Crops (pp. 518–549). Academic Press. 10.1016/B978-0-12-819194-1.00017-7

Langfelder, P., & Horvath, S. (2008). WGCNA: an R package for weighted correlation network analysis. BMC Bioinformatics, 9, 559. 10.1186/1471-2105-9-559

Li, L., Tian, Z., Chen, J., Tan, Z., Zhang, Y., Zhao, H., Wu, X., Yao, X., Wen, W., Chen, W., & Guo, L. (2023). Characterization of novel loci controlling seed oil content in Brassica napus by marker metabolite-based multi-omics analysis. Genome Biol, 24(1), 141. 10.1186/s13059-023-02984-z

Li, Z., Costamagna, A. C., Beran, F., & You, M. (2024). Biology, Ecology, and Management of Flea Beetles in Brassica Crops. Annu Rev Entomol, 69, 199–217. 10.1146/annurev-ento-033023-015753

Llanes, A., Andrade, A., Alemano, S., & Luna, V. (2018). Chapter 6 - Metabolomic Approach to Understand Plant Adaptations to Water and Salt Stress. In P. Ahmad, M. A. Ahanger, V. P. Singh, D. K. Tripathi, P. Alam, & M. N. Alyemeni (Eds.), Plant Metabolites and Regulation Under Environmental Stress (pp. 133–144). Academic Press. 10.1016/B978-0-12-812689-9.00006-6

Martins, M. C. M., Mafra, V., Monte-Bello, C. C., & Caldana, C. (2021). The Contribution of Metabolomics to Systems Biology: Current Applications Bridging Genotype and Phenotype in Plant Science. In F. Vischi Winck (Ed.), Advances in Plant Omics and Systems Biology Approaches (pp. 91–105). Springer International Publishing. 10.1007/978-3-030-80352-0_5

Matthees, H. L., Thom, M. D., Gesch, R. W., & Forcella, F. (2018). Salinity tolerance of germinating alternative oilseeds. Industrial Crops and Products, 113, 358–367. 10.1016/j.indcrop.2018.01.042

Nunn, A., Rodriguez-Arevalo, I., Tandukar, Z., Frels, K., Contreras-Garrido, A., Carbonell-Bejerano, P., Zhang, P., Ramos Cruz, D., Jandrasits, K., Lanz, C., Brusa, A., Mirouze, M., Dorn, K., Galbraith, D. W., Jarvis, B. A., Sedbrook, J. C., Wyse, D. L., Otto, C., Langenberger, D., . . . Chopra, R. (2022). Chromosome-level Thlaspi arvense genome provides new tools for translational research and for a newly domesticated cash cover crop of the cooler climates. Plant Biotechnol J, 20(5), 944–963. 10.1111/pbi.13775

Onyilagha, J., Bala, A., Hallett, R., Gruber, M., Soroka, J., & Westcott, N. (2003). Leaf flavonoids of the cruciferous species, Camelina sativa, Crambe spp., Thlaspi arvense and several other genera of the family Brassicaceae. Biochemical Systematics and Ecology, 31(11), 1309–1322. 10.1016/S0305-1978(03)00074-7

Onyilagha, J. C., Gruber, M. Y., Hallett, R. H., Holowachuk, J., Buckner, A., & Soroka, J. J. (2012). Constitutive flavonoids deter flea beetle insect feeding in Camelina sativa L. Biochemical Systematics and Ecology, Volume 42, 128–133. 10.1016/j.bse.2011.12.021

Prieto, M. A., López, C. J., & Simal-Gandara, J. (2019). Chapter Six - Glucosinolates: Molecular structure, breakdown, genetic, bioavailability, properties and healthy and adverse effects. In I. C. F. R. Ferreira & L. Barros (Eds.), Advances in Food and Nutrition Research (Vol. 90, pp. 305–350). Academic Press. 10.1016/bs.afnr.2019.02.008

Qi, J., Li, K., Shi, Y., Li, Y., Dong, L., Liu, L., Li, M., Ren, H., Liu, X., Fang, C., & Luo, J. (2021). Cross-Species Comparison of Metabolomics to Decipher the Metabolic Diversity in Ten Fruits. Metabolites, 11(3). 10.3390/metabo11030164

Qi, J., Wei, J., Liao, D., Ding, Z., Yao, X., Sun, P., & Li, X. (2022). Untargeted Metabolomics Analysis Revealed the Major Metabolites in the Seeds of four Polygonatum Species. Molecules, 27(4). 10.3390/molecules27041445

Ramaroson, M. L., Koutouan, C., Helesbeux, J. J., Le Clerc, V., Hamama, L., Geoffriau, E., & Briard, M. (2022). Role of Phenylpropanoids and Flavonoids in Plant Resistance to Pests and Diseases. Molecules, 27(23). 10.3390/molecules27238371

Raza, A., Hafeez, M. B., Zahra, N., Shaukat, K., Umbreen, S., Tabassum, J., Charagh, S., Khan, R. S. A., & Hasanuzzaman, M. (2020). The Plant Family Brassicaceae: Introduction, Biology, And Importance. In M. Hasanuzzaman (Ed.), The Plant Family Brassicaceae: Biology and Physiological Responses to Environmental Stresses (pp. 1–43). Springer Singapore. 10.1007/978-981-15-6345-4_1

Salam, U., Ullah, S., Tang, Z. H., Elateeq, A. A., Khan, Y., Khan, J., Khan, A., & Ali, S. (2023). Plant Metabolomics: An Overview of the Role of Primary and Secondary Metabolites against Different Environmental Stress Factors. Life (Basel), 13(3). 10.3390/life13030706

Seifi, H. S., & Shelp, B. J. (2019). Spermine Differentially Refines Plant Defense Responses Against Biotic and Abiotic Stresses. Front Plant Sci, 10, 117. 10.3389/fpls.2019.00117

Shao, J., Huang, K., Batool, M., Idrees, F., Afzal, R., Haroon, M., Noushahi, H. A., Wu, W., Hu, Q., Lu, X., Huang, G., Aamer, M., Hassan, M. U., & El Sabagh, A. (2022). Versatile roles of polyamines in improving abiotic stress tolerance of plants. Front Plant Sci, 13, 1003155. 10.3389/fpls.2022.1003155

Sharma, A., Shahzad, B., Rehman, A., Bhardwaj, R., Landi, M., & Zheng, B. (2019). Response of Phenylpropanoid Pathway and the Role of Polyphenols in Plants under Abiotic Stress. Molecules, 24(13). 10.3390/molecules24132452

Soroka, J., & Grenkow, L. (2013). Susceptibility of brassicaceous plants to feeding by flea beetles, Phyllotreta spp. (Coleoptera: Chrysomelidae). J Econ Entomol, 106(6), 2557–2567. 10.1603/ec13102

Variyar, P. S., Banerjee, A., Akkarakaran, J. J., & Suprasanna, P. (2014). Chapter 12 - Role of Glucosinolates in Plant Stress Tolerance. In P. Ahmad & S. Rasool (Eds.), Emerging Technologies and Management of Crop Stress Tolerance (pp. 271–291). Academic Press. 10.1016/B978-0-12-800876-8.00012-6

Vogt, T. (2010). Phenylpropanoid biosynthesis. Mol Plant, 3(1), 2–20. 10.1093/mp/ssp106

Vollmann, J., & Eynck, C. (2015). Camelina as a sustainable oilseed crop: contributions of plant breeding and genetic engineering. Biotechnol J, 10(4), 525–535. 10.1002/biot.201400200

Wang, R., Ren, C., Dong, S., Chen, C., Xian, B., Wu, Q., Wang, J., Pei, J., & Chen, J. (2021). Integrated Metabolomics and Transcriptome Analysis of Flavonoid Biosynthesis in Safflower (Carthamus tinctorius L.) With Different Colors. Front Plant Sci, 12, 712038. 10.3389/fpls.2021.712038

Wang, S., Alseekh, S., Fernie, A. R., & Luo, J. (2019). The Structure and Function of Major Plant Metabolite Modifications. Mol Plant, 12(7), 899–919. 10.1016/j.molp.2019.06.001

Wang, S., Tu, H., Wan, J., Chen, W., Liu, X., Luo, J., Xu, J., & Zhang, H. (2016). Spatio-temporal distribution and natural variation of metabolites in citrus fruits. Food Chem, 199, 8–17. 10.1016/j.foodchem.2015.11.113

Yan, S., Bhawal, R., Yin, Z., Thannhauser, T. W., & Zhang, S. (2022). Recent advances in proteomics and metabolomics in plants. Mol Hortic, 2(1), 17. 10.1186/s43897-022-00038-9

Yonekura-Sakakibara, K., & Saito, K. (2009). Functional genomics for plant natural product biosynthesis. Nat Prod Rep, 26(11), 1466–1487. 10.1039/b817077k

Zanetti, F., Isbell, T. A., Gesch, R. W., Evangelista, R. L., Alexopoulou, E., Moser, B., & Monti, A. (2019). Turning a burden into an opportunity: Pennycress (Thlaspi arvense L.) a new oilseed crop for biofuel production. Biomass and Bioenergy, 130, 105354. 10.1016/j.biombioe.2019.105354

Zhao, S., Li, H., Han, W., Chan, W., & Li, L. (2019). Metabolomic Coverage of Chemical-Group-Submetabolome Analysis: Group Classification and Four-Channel Chemical Isotope Labeling LC-MS. Analytical Chemistry, 91(18), 12108–12115. 10.1021/acs.analchem.9b03431

Zhao, Y., Zhou, M., Xu, K., Li, J., Li, S., Zhang, S., & Yang, X. (2019). Integrated transcriptomics and metabolomics analyses provide insights into cold stress response in wheat. The Crop Journal, 7(6), 857–866. 10.1016/j.cj.2019.09.002

